# Improved Zika virus plaque assay using Vero/TMPRSS2 cell line

**DOI:** 10.1101/2024.08.16.608323

**Authors:** Olukunle O. Oluwasemowo, Monica E. Graham, Deepa K. Murugesh, Monica K. Borucki

**Affiliations:** Biosciences and Biotechnology Division, Physical and Life Sciences Directorate, Lawrence Livermore National Laboratory, Livermore, CA 94550, USA

## Abstract

Plaque assay is the gold standard for the quantification of viable cytopathic viruses like Zika virus (ZIKV). Some strains of ZIKV produce plaques that are very difficult to accurately visualize and count on the commonly used Vero cell line. From data generated in our lab, we became curious if Vero/TMPRSS2 cells may be a better alternative, therefore we compared the plaque forming units (PFU) of two strains of ZIKV on Vero/TMPRSS2 cells to those produced by Vero cells. We also compared the virus stock titer generated on Vero/TMPRSS2 cells to that generated by the Vero cell line. Although Vero cells generated higher quantity of ZIKV stocks, Vero/TMPRSS2 cells produced plaques with significantly improved morphology and visibility and may therefore be a better alternative to use for performing plaque assays for strains of ZIKV that are more difficult to titer on regular Vero cells.

## IMPORTANCE

While there are several methods of viral quantification, plaque assay remains the gold standard for accurate quantification of replication-competent, cytopathic viruses including Zika virus (ZIKV). Vero cells are commonly used to titer ZIKV stock via plaque assay. Prior to the initiation of this study, we observed that ZIKV-PRV strain plaque assays using Vero cells often yielded plaques that were very small, overly diffuse, and hard to count. We also observed that the ZIKV-CAM strain often did not produce plaques at all on Vero cells. This study shows that Vero cells expressing TMPRSS2 improve the morphology and visibility of plaques produced by ZIKV-PRV and ZIKV-CAM compared to regular Vero cells and may therefore be a better alternative to use for performing plaque assays for these strains and perhaps for other strains of ZIKV that are difficult to titer. However, Vero cells proved to be superior for generating high titer stock.

## OBSERVATION

We observed that for both strains of ZIKV tested, the plaques from the Vero cells were diffuse, indiscrete, and very difficult to enumerate regardless of the overlay used and the scientist that performed the assay (Fig 1). However, the plaques of both strains were clearly visible and distinct enough to be counted with ease when plaqued on Vero/TMPRSS2 cells.

**Fig 1.**
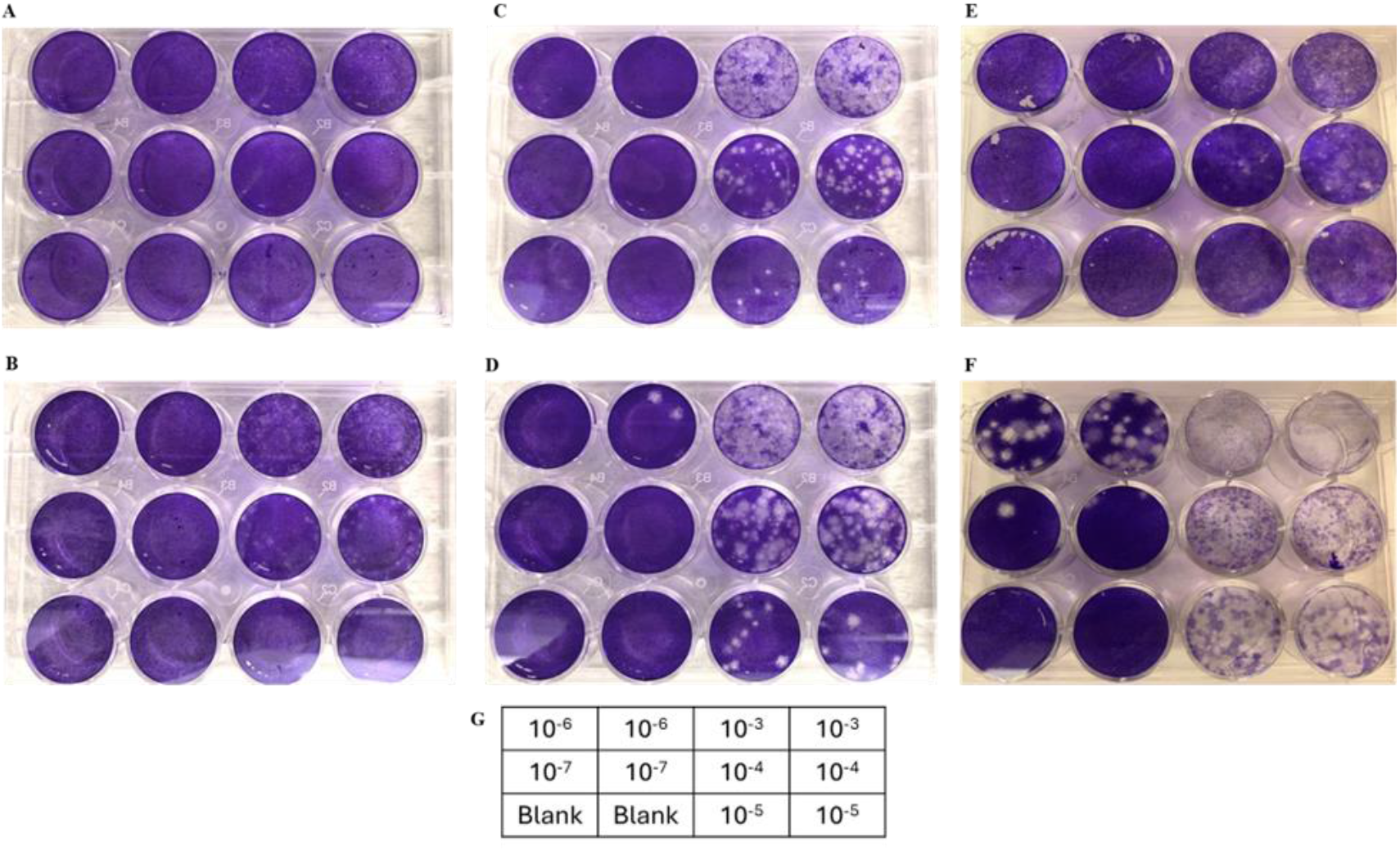
Plaque morphologies of the ZIKV PRV and CAM on Vero and Vero/TMPRSS2 cells. ZIKV-PRV on Vero cells at (A) 4 days post-infection (dpi) and (B) 5 dpi. ZIKV-PRV on Vero/TMPRSS2 cells at (C) 4 dpi and (D) 5 dpi. ZIKV-CAM on (E) Vero cells and (F) Vero/TMPRSS2 at 5 dpi. The plate map (G) shows the dilution factor of the inoculum.

Following the observation of improved plaque visualization and enumeration in assays performed with Vero/TMPRSS2 cells, we were curious to know if Vero/TMPRSS2 cells could also produce higher titer of virus than Vero cells when used for ZIKV propagation. Virus stocks were generated on both cell lines and titered by TCID_50_ on Vero/TMPRSS2 cells. In Vero/TMPRSS2 cells, cytopathic effect (CPE) began 3 dpi, with significant 50-70% CPE occurring 4 dpi. Significant loss of the cell monolayer, over 90%, continued through day 7. Viral titer for both ZIKV-CAM and ZIKV-PRV peaked at 3 dpi and decreased by 100-fold on day 4 post-infection, with another 10-fold decrease on day 5 (Fig 2).

**Fig 2.**
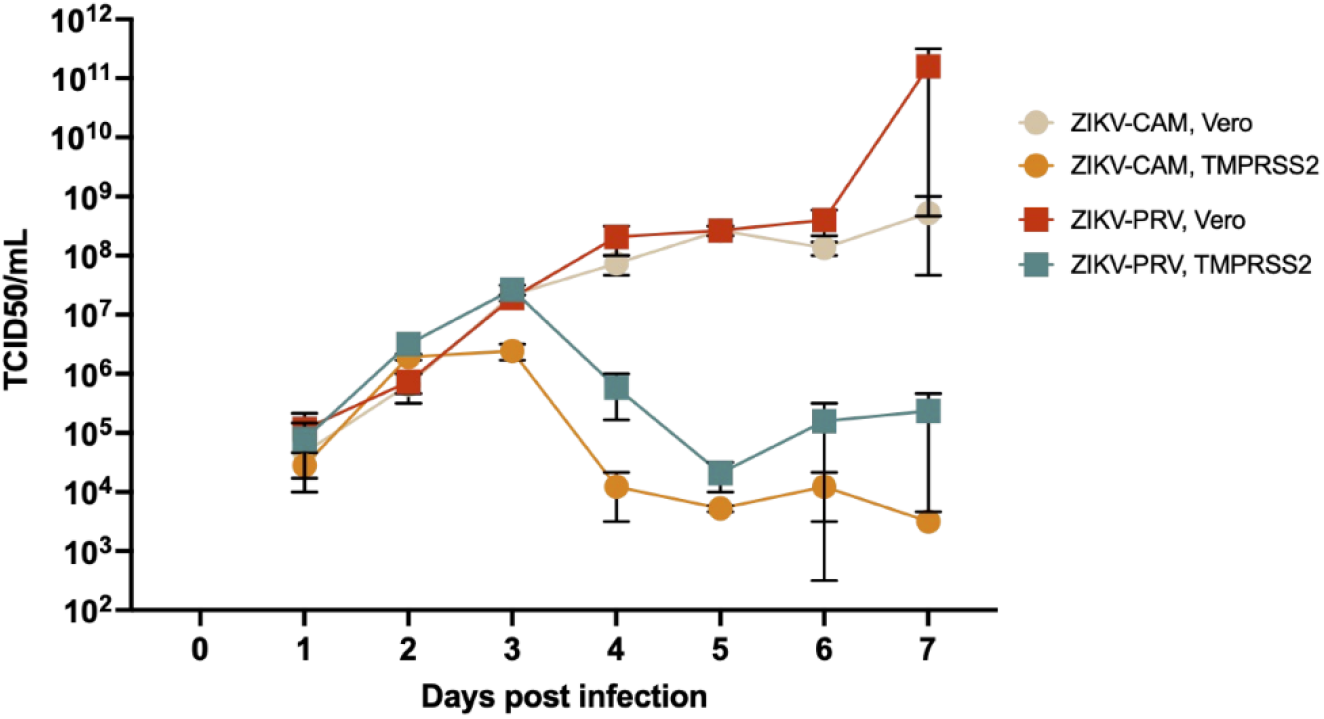
Titer of Zika virus on Vero and Vero/TMPRSS2 cells.

In Vero cells, significant CPE was not observed following ZIKV-CAM or ZIKV-PRV infection out to 10 dpi. However, viral titer for both strains steadily increased during the time course, peaking at 7 dpi. ZIKV-CAM growth in Vero cells peaked at 100-fold higher titer compared to growth in Vero/TMPRSS2 cells. Similarly, ZIKV-PRV growth in Vero cells peaked at 10,000-fold higher titer compared to growth in Vero/TMPRSS2 cells.

## Discussion

We observed that Vero/TMPRSS2 cells produced plaques with significantly improved morphology and visibility, and this may enable more strains of ZIKV to be titered reliably. Since these data showed an improvement in cytopathic effect in Vero/TMPRSS2 cells via plaque assay, we hypothesized that virus propagation would also be improved in TMPRSS2-expressing cells. For ZIKV-PRV and ZIKV-CAM propagated in Vero/TMPRSS2 cells we observed peak virus titer of approximately 10^7^ and 10^6^ TCID_50_/mL respectively at 3 days post inoculation. Viral titer continued to increase in Vero cells unlike the 3 days post infection peak observed in Vero/TMPRSS2 cells. At 7 days post infection, when the peak of virus replication was observed in Vero cells, the virus titer of 10^11^ and 10^8^ TCID_50_/mL for ZIKV-PRV and ZIKV-CAM, respectively, were significantly higher than the peak titer in the Vero/TMPRSS2 cells. This clearly indicates that the inability of ZIKV to form distinctive plaques on Vero cells did not impair the effective viral replication, and that Vero cell remains the cell of choice to employ where there is the need to maximize ZIKV titer. A potential mechanism for rapid virus production and defined plaque formation is that TMPRSS2 helps facilitate viral maturation by cleaving prM on immature or partially mature ZIKV virions at the furin cleavage site, allowing for more virus to enter more cells. This is supported by the early rapid growth and CPE caused by ZIKV observed in our plaque assay and viral propagation data. For our plaque assays an increase in mature virions caused by TMPRSS2 cleavage would allow for increased viral entry and CPE of cells, thus allowing plaques to be visualized more clearly in a shorter time frame compared to standard Vero cells. Since this type of titer assay is done over a short period of time and viral spread is limited by a viscous overlay, widespread CPE is not a concern for negatively affecting the results. However, for viral propagation this increase in mature virions is likely killing too many cells too quickly to sustain high titer virus production.

In conclusion, the results from our study demonstrated the utility of Vero/TMPRSS2 cells for generating distinct and easy-to-count plaques for strains of ZIKV as compared to the regular Vero cell line. However, Vero cells proved to be better for generating high titered ZIKV stock.

## Methods

### Cells and Viruses

The Vero and Vero/TMPRS2 cells used in this study were of similar level of low passage and monolayers were of similar confluency when infected. Vero cells (P5) (ATCC^®^ CCL-81™) and Vero/TMPRSS2 cells (P6) were grown at 37°C in media Dulbecco’s Modified Eagle Medium (DMEM) supplemented with 10% heat-inactivated fetal bovine serum FBS (Gibco), 1% penicillin/streptomycin, 1% non-essential amino acids, and 1% sodium pyruvate. Puerto Rico PRVABC59/2015 and Cambodian (FSS13025) strains of ZIKV with approximately 1.5 × 10^5 TCID50/mL and 1.5 × 10^5 TCID50/mL respectively were used for the assay.

### Plaque Assay

The Vero or Vero/TMPRSS2 cells were seeded into 12-well plates at a density of 2e^5^ cells/well in 1mL of 1X DMEM with 10% FBS and 1% penicillin-streptomycin and infected 24 hours post seeding at a confluency of 80%. The cells were infected with 200μL of tenfold serially diluted viruses and incubated for one hour, with intermittent rocking. Post-infection, the cells were overlaid with 1mL of semisolid Microcrystaline Cellulose (MCC) media (50% 2X MEM with 8% FBS, 25% 2.4% MCC, and 25% sterile tissue culture water). Plates were incubated for 4 to 5 days at 37°C. Plaques were counted after removing the overlay, washing once with 1X PBS, and staining with 0.25% (w/v) crystal violet in 100% methanol for 10 minutes.

### Virus Stock Generation Assay

Vero or Vero/TMPRSS2 cells were seeded into tissue-culture treated T-75 flasks at a density of 6.7e^6^ cells/flask. ZIKV-PRV and ZIKV-CAM were prepared to infect the cells at an MOI of 0.01 TCID_50_/mL and 0.01 PFU/mL respectively. Cells were inoculated with respective virus and incubated for one hour at 37°C, with intermittent flask rocking. Viral inoculum was removed, and cells were washed three times with 5 mL 1X PBS. Ten milliliters of media was added to each flask. Supernatant samples were collected from each flask (day 0 sample) for downstream analysis and stored at −80°C in duplicate aliquot of 500μL. After the day 0 sample collection, 1mL of fresh viral media was added to each flask.

## ACKNOWLEDGMENTS

Vero-TMPRSS2 cells were received from Sean Whelan’s laboratory (Univ. Washington). This work was performed under the auspices of the U.S. Department of Energy by Lawrence Livermore National Laboratory under Contract DE-AC52-07NA27344.

## Notes

### Competing Interest Statement

The authors have declared no competing interest.

